# Late-onset glaucoma in *Yap* conditional knockout mouse

**DOI:** 10.1101/2022.05.16.492143

**Authors:** Juliette Bitard, Elodie-Kim Grellier, Sophie Lourdel, Helena Prior Filipe, Annaïg Hamon, François Fenaille, Florence Anne Castelli, Emeline Chu-Van, Jérôme E. Roger, Morgane Locker, Muriel Perron

**Affiliations:** Paris-Saclay Institute of Neuroscience, Université Paris-Saclay, CNRS, CERTO-Retina France, 91400 Saclay, France; West Lisbon Hospitals Center. Hospital Egas Moniz; Center for International Investigation Egas Moniz; Université de Rennes Inserm, EHESP, Irset (Institut de recherche en santé, environnement et travail) - UMR_S 1085, F-35000 Rennes, France; Université Paris-Saclay, CEA, INRAE, Département Médicaments et Technologies pour la Santé (DMTS), MetaboHUB, F-91191 Gif sur Yvette, France

**Keywords:** YAP, Ciliary body, Müller cells, Glaucoma

## Abstract

Glaucoma is an optic neuropathy often referred to as “the silent thief of sight”, due to its late diagnosis, which is generally made when degeneration of the optic nerve and retinal ganglion cells is already well under way. It is thus of utmost importance to have a better understanding of the disease, and to investigate more deeply the early causes of glaucoma. The transcriptional coactivator YAP recently emerged as an important regulator of eye homeostasis and is drawing attention in the glaucoma research field. Here we show that *Yap* conditional knockout mice (*Yap* cKO), in which the deletion of *Yap* is induced in both Müller glia (*i.e*. the only retinal YAP-expressing cells) and the non-pigmented epithelial cells of the ciliary body, exhibit breakdown of the aqueous-blood barrier accompanied by progressive collapse of the ciliary body as we observed in human uveitic patients. In addition, aged *Yap* cKO mice harbor glaucoma features, including alteration of glutamate recycling, deregulation of key homeostatic Müller-derived proteins, retinal vascular defects, optic nerve degeneration, and retinal ganglion cell death. Together, our findings reveal the essential role of YAP in preserving the ciliary body and the retinal ganglion cells, thereby preventing the onset of glaucoma features.

## INTRODUCTION

Glaucoma is a multifactorial neuropathy characterized by optic nerve damage and progressive loss of retinal ganglion cells (RGCs). The etiologic diversity of the disease, which leads to more than ten clinical subtypes (*e*.*g*. open-angle glaucoma, normal tension glaucoma, uveitic glaucoma, pigmentary glaucoma, etc), as well as the very limited number of animal models, have hindered our understanding of the underlying pathological mechanisms. Dysfunctions of both the anterior chamber and the neuroretina take part to the development of glaucoma. In the anterior chamber, inflammation of the uvea (*i*.*e*. iris, choroid, ciliary body) called uveitis, pigment dispersion from the iris, and increased intraocular pressure (IOP), constitute well-known risk factors. High IOP results from the imbalance between production of the aqueous humor by the non-pigmented epithelium (NPE) of the ciliary body, and its drainage by the trabecular meshwork in the iridocorneal angle ^1^. Since IOP is the only modifiable factor in glaucoma, it constitutes the target of first-line medications. Nevertheless, between 30% to 90% of glaucomatous patients will not present this trait, suggesting the existence of other pathogenic processes. In line with this, both normal and high-tension glaucoma are accompanied at the neuroretinal level, by chronic gliosis, oxidative stress, and excitotoxicity ^2^. These pathological features might be explained, at least in part, by Müller glial cell dysfunction ^3–5^. Among the fundamental roles of Müller cells is the regulation of extracellular glutamate levels through its recycling into glutamine that further fuels neurons ^6^. In glaucomatous patients and in several experimental models of glaucoma, an increase in extracellular glutamate levels was reported to contribute to RGC death ^7^.

Despite major breakthroughs in the global understanding of glaucoma, new animal models are still needed to further dissect the complex etiology of this group of diseases. The Hippo pathway effector YAP recently emerged as an orchestrator of eye homeostasis ^8^ and we showed in particular that its haploinsufficiency in 12-month-old mice results in photoreceptor degeneration ^9^. With regards to the glaucoma research field, a genome wide association study identified *Yap* as a new risk locus for primary open-angle glaucoma ^10^. In line with this, YAP is expressed in several ocular cell types, ***including those affected in glaucoma (i*.*e. the NPE of the ciliary body, the trabecular*** meshwork, and Müller cells) ^11^. Functional studies revealed its contribution to human trabecular meshwork cell stiffening *in vitro*, through its mechanotransducer function ^12,13^. This further substantiates the idea that YAP dysfunction might play a role in glaucoma since changes in mechanical trabecular meshwork properties are proposed to underlie the increased resistance to aqueous humor outflow, and thereby to participate in the IOP raise observed in glaucomatous patients ^14^. Whether altered *Yap* expression in NPE cells of the ciliary body or in Müller glial cells could also contribute to glaucoma development has not been investigated yet. Here, taking advantage of *Yap* conditional knockout mice that we previously described ^15^, we found that selective loss of YAP in Müller cells and in the ciliary body triggers pathogenic features of secondary open-angle glaucoma. Our study reveals in particular that YAP acts as an essential player in ciliary body maintenance and Müller cells homeostasis, thereby suggesting that deregulation of this cofactor may impact glaucoma progression in human.

## RESULTS

### The ciliary body progressively collapses with age in Yap cKO mice

In order to evaluate the impact of *Yap* deletion in Müller cells and ciliary body, we used *Yap*^flox/flox^;Rax-Cre^ERT2^ mice (*Yap* cKO) that we previously generated ^15^. In the postnatal or adult eye, Cre recombinase expression pattern is restricted to Müller glia and NPE cells of the ciliary body where YAP was previously shown to be expressed ^15,16^. *Yap* deletion was induced at P10, when both Müller and NPE cells are already differentiated ^17,18^. We first assessed ciliary body morphology during aging. Two months after *Yap* deletion induction, around 60% of the animals exhibited distended ciliary processes indicative of cell-cell contact loss (Figure 1A). The ciliary processes thereafter further regressed, with complete collapse progressively occurring between six and nine months of age without angle closure (Figure 1A, 1B). Confirming its dramatic disorganization, ciliary bodies of 9-month-old *Yap* cKO mice exhibited an increased surface compared to controls ones (Figure 1C). To our knowledge, this latter feature has been described only in uveitic patients ^19^. We next sought to further characterize the ciliary body defects of uveitic patients by performing ultrasound biomicroscopy (UBM) examination. Very interestingly, UBM images revealed distension (low IOP patients) or shortening (high IOP patients) of ciliary body processes (Figure 1D) as observed in *Yap* cKO mice.

**Figure 1.**
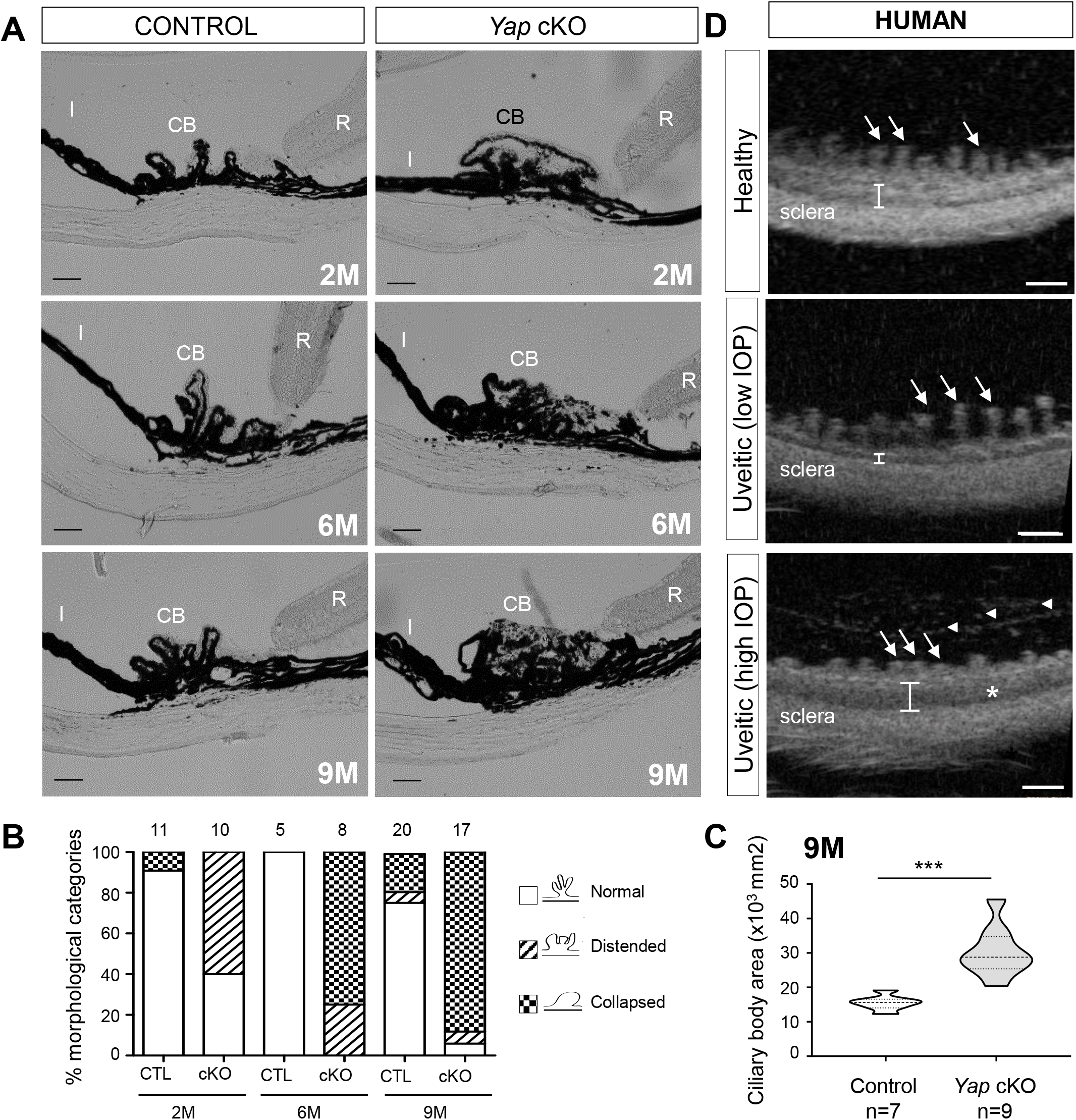
The ciliary body of *Yap* cKO mice progressively collapses as in human uveitic patients. (A)Bright field micrographs from 2-, 6-, 9-month-old (2M, 6M, 9M) control (*Yap*^*flox/flox*^) and *Yap* cKO (*Yap*^*flox/flox*^*;RaxCre*^*ERT2*^) ciliary body sections. CB: Ciliary body, I: Iris, R: Retina. Scale bars: 50 mm. (B) Quantification of control (CTL) and *Yap* cKO eyes harboring normal or defective ciliary bodies. We defined the ciliary body morphology according to the shape of its processes, which were either “distended” with loss of clearly defined separation between the non-pigmented and the pigmented epithelium, or “collapsed” when the processes had completely disappeared. The number of individual eyes analyzed for each condition is indicated above the stacked histograms. (C) Quantification of ciliary body area in 9-month-old (9M) control and *Yap* cKO individuals. In these violin-plots, the thick dotted line represents the median, and the thin dotted lines the interquartiles. The number of analyzed ciliary bodies in each condition is indicated below the x-axis. Statistics: Mann-Whitney test, ***p≤ 0.001. (D) Transverse section images from human healthy and chronic uveitic patients taken with an ultrasound biomicroscope. Compared to the healthy individual, the uveitic patient presenting with low IOP shows atrophic (almost absent) pars plicata, thinner ciliary muscle area (white bracket), and elongated ciliary processes (arrows). The second uveitic patient with high IOP presents with pars plicata oedema (asterisk), increased thickness of ciliary muscle area (bracket), and shortened ciliary processes (arrows). White arrowheads point to putative inflammatory cells and fibrin deposition Scale bars: 1 mm.

Overall, these data demonstrate that YAP is required for the structural maintenance of the ciliary body. They also interestingly highlight a phenotypic resemblance between *Yap* cKO mice and uveitic patients.

### Yap cKO mice show alterations of the iridal blood-aqueous-barrier

We next sought for other signs of uveitis in *Yap* cKO eyes, such as a breakdown of the blood-aqueous-barrier (BaB). As evidenced by albumin immunostaining ^20^, we found indeed that 100% of *Yap* cKO eyes exhibited iridal blood leakage starting by two months of age (Figure 2A). In aged *Yap* cKO eyes, IsoB4-labelled iridal blood vessel walls showed signs of degeneration compared to control ones, appearing either dilated or degraded. In addition, proteinaceous exudates likely emanating from the iridal endothelial cell layer were observed (Figure 2B). To confirm BaB disruption at the level of the iris, we performed fluorescein angiography on 9-month-old mice (Figure 2C). Compared with control eyes, where iridal vascularization consists in thin regular blood vessels, *Yap* cKO eyes exhibited thicker blood vessels, highly visible at the arcade of the pupillary margin and covering a larger vascular area. They also displayed blood leakage as indicated by iridal zones presenting diffuse fluorescein staining.

**Figure 2.**
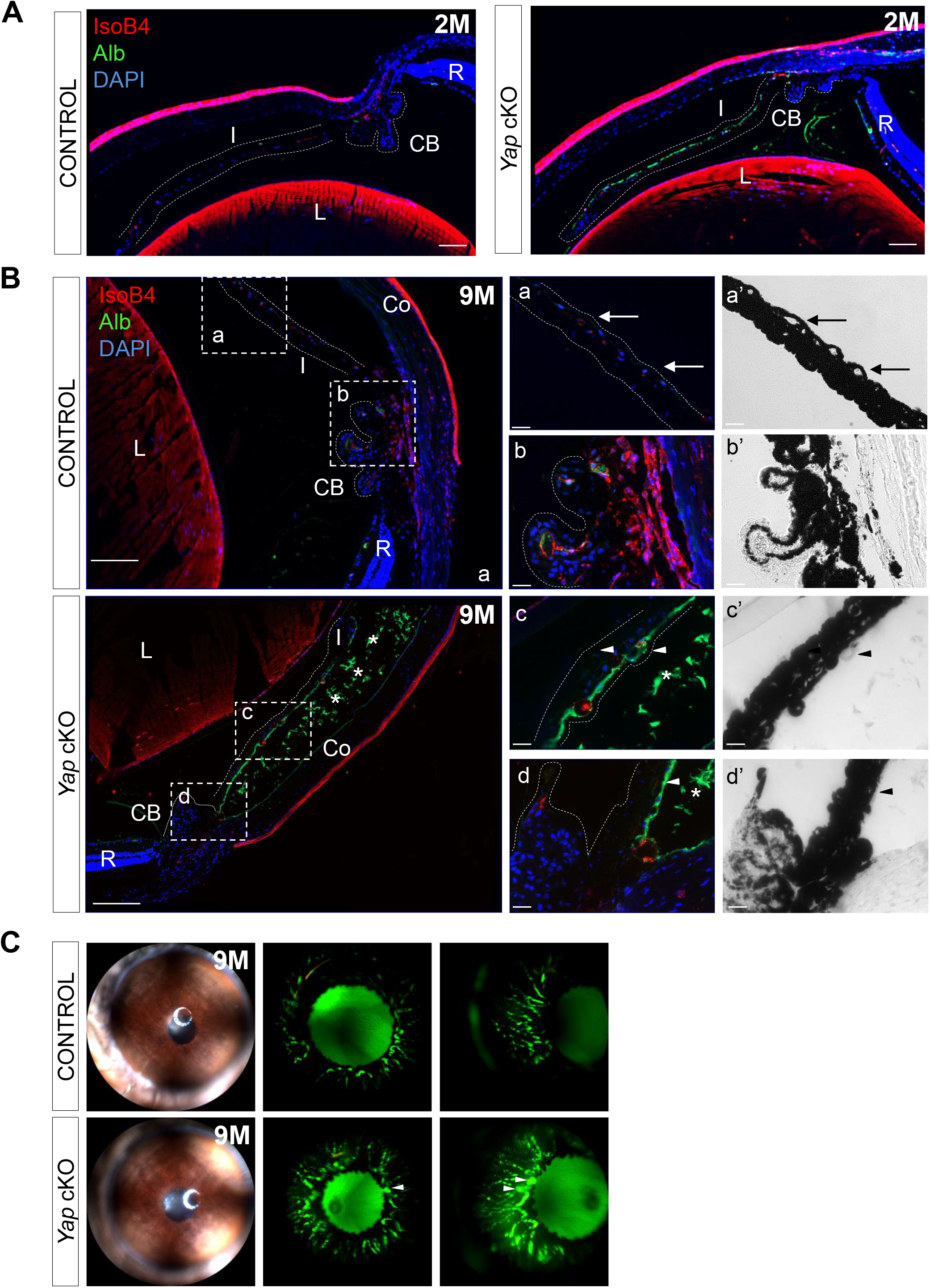
*Yap* cKO eyes exhibit breakdown of the blood-aqueous-barrier. (A, B) Eye sections from control (*Yap*^*flox/flox*^) and *Yap* cKO (*Yap*^*flox/flox*^*;RaxCre*^*ERT2*^) eyes at 2 (2M, n=4; A) or 9 months of age (9M, n=4; B). Sections were stained with isolectin B4 (red) to assess blood vessel wall integrity, and with anti-albumin antibody (green) to assess blood leakage from capillaries. Nuclei were counterstained with DAPI (blue). Dashed white lines delineate the iris and the ciliary body. Regions inside the dashed white squares are depicted at a higher magnification on the right panels. Shown are either fluorescence (a, b, c, d) or brightfield (a’, b’, c’, d’) images. Asterisks indicate proteinaceous exudates, arrows point to normal iridal blood vessels, arrowheads indicate abnormal blood vessels. CB: Ciliary body, Co: Cornea, I: Iris, L: Lens, R: Retina. At least 3 individuals per condition were analyzed. Scale bars: left panels, 100 mm; right panels, 20 mm. (C) Fundus and fluorescein angiography (en-face and profile views) of 9-month-old (9M) control (n=5) and *Yap* cKO (n=7) iris. Arrowheads indicate microaneurisms.**L**

Taken together, our results demonstrate that loss of blood vessel integrity in the iris constitutes an early trait of *Yap* cKO eyes.

### Yap cKO mice harbor glaucoma features

The BaB breakdown and ciliary body collapse occurring in *Yap* cKO mice are reminiscent of the phenotype reported in the widely-used secondary glaucoma mouse model DBA/2J ^21,22^. This led us to hypothesize that *Yap* cKO individuals might also develop glaucomatous features with age. Considering the essential role of NPE in aqueous humor production, we examined whether the collapsed ciliary body of *Yap* cKO mice might be associated with abnormal IOP. We however did not detect any differences in IOP values between control and *Yap* cKO eyes whatever their age (Figure 3A).

**Figure 3.**
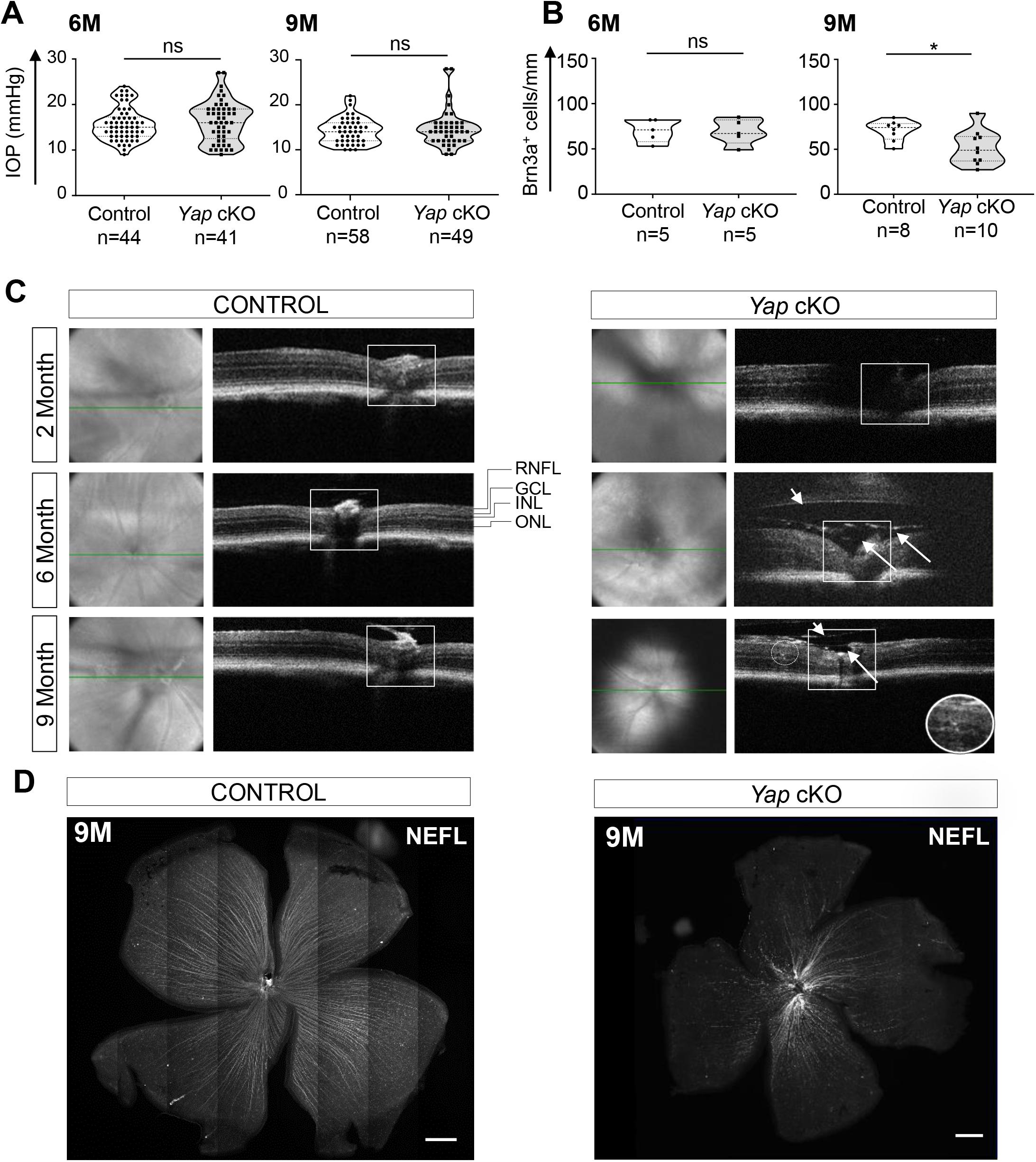
*Yap* cKO mice harbor glaucoma features. (A)Measurement of intraocular pressure (IOP) on 6-or 9-month-old (6M and 9M) control (*Yap*^*flox/flox*^) and *Yap* cKO (*Yap*^*flox/flox*^*;RaxCre*^*ERT2*^) mice. (B) Quantification of Brn3a-positive cells in the ganglion cell layer (GCL) of control (*Yap*^*flox/flox*^) and *Yap* cKO (*Yap*^*flox/flox*^*;RaxCre*^*ERT2*^) retinas at 6 and 9 months of age (6M, 9M). In violin plots (A, B), the thick dotted line represents the median and the thin dotted lines the interquartiles. The number of analyzed eyes in each condition is indicated below the x-axis. Statistics: Mann-Whitney test, ns: non-significant, *p≤ 0.05. (C) Representative OCT B-scan (right) and fundus (left) images of 2-, 6-, and 9-month-old (2M, 6M, 9M) control (*Yap*^*flox/flox*^) and *Yap* cKO (*Yap*^*flox/flox*^*;RaxCre*^*ERT2*^) retinas. The optic nerve head excavation is delineated by a white square. In *Yap* cKO eyes, long white arrows point to oedema, while short ones indicate vitreal detachment. A region containing intra-retinal cysts in the INL is surrounded by a white circle and enlarged in the inset. At least 3 individuals per condition were analyzed. GCL: ganglion cell layer, INL: inner nuclear layer, ONL: outer nuclear layer, RNFL: retinal nerve fiber layer. (D) Representative flat-mounted 9-month-old control (n=3) and *Yap* cKO (n=5) retinas immunostained with anti-light chain neurofilament antibody. Scale bars: 500 µm.

We next further examined potential glaucomatous features in *Yap* cKO eyes by assessing whether RGCs might be affected. At 6 months of age, the proportion of RGCs within the ganglion cell layer was unchanged compared to control retinas. However, it was significantly decreased by 47% in 9-month-old individuals (Figure 3B). We then turned to optic coherence tomography (OCT) to visualize the optic nerve head (ONH). Control mice harbored a uniformly flat ONH whatever their age. In contrast, *Yap* cKO ONH presented with a typical deformation called “cupping” from 2 months onwards (Figure 3C). In addition, aged *Yap* cKO eyes harbored edema inside or near the ONH, as well as occasional vitreous detachment and cysts near the ONH (Figure 3C). Finally, we found that these ONH defects were accompanied by nerve fiber degeneration, as assessed by immunostaining of light chain neurofilament on flat mounted 9-month-old retinas (Figure 3D).

Collectively, these data indicate that RGC integrity and survival are affected in the neuroretina of 9-month-old *Yap* cKO mice, a typical feature of glaucoma. Furthermore, we show that ONH cupping is an early event that precedes RGC loss.

### Reactive gliosis and retinal vasculature defects precede RGCs death in Yap cKO retinas

Müller cell gliosis is a hallmark of glaucoma and has been shown to participate in its physiopathology ^23,24^. Analysis of GFAP expression by both western-blot and immunostaining revealed onset of reactive gliosis in *Yap* cKO mice from 6 months of age. This further worsened in older animals (Figure 4A and 4B).

**Figure 4.**
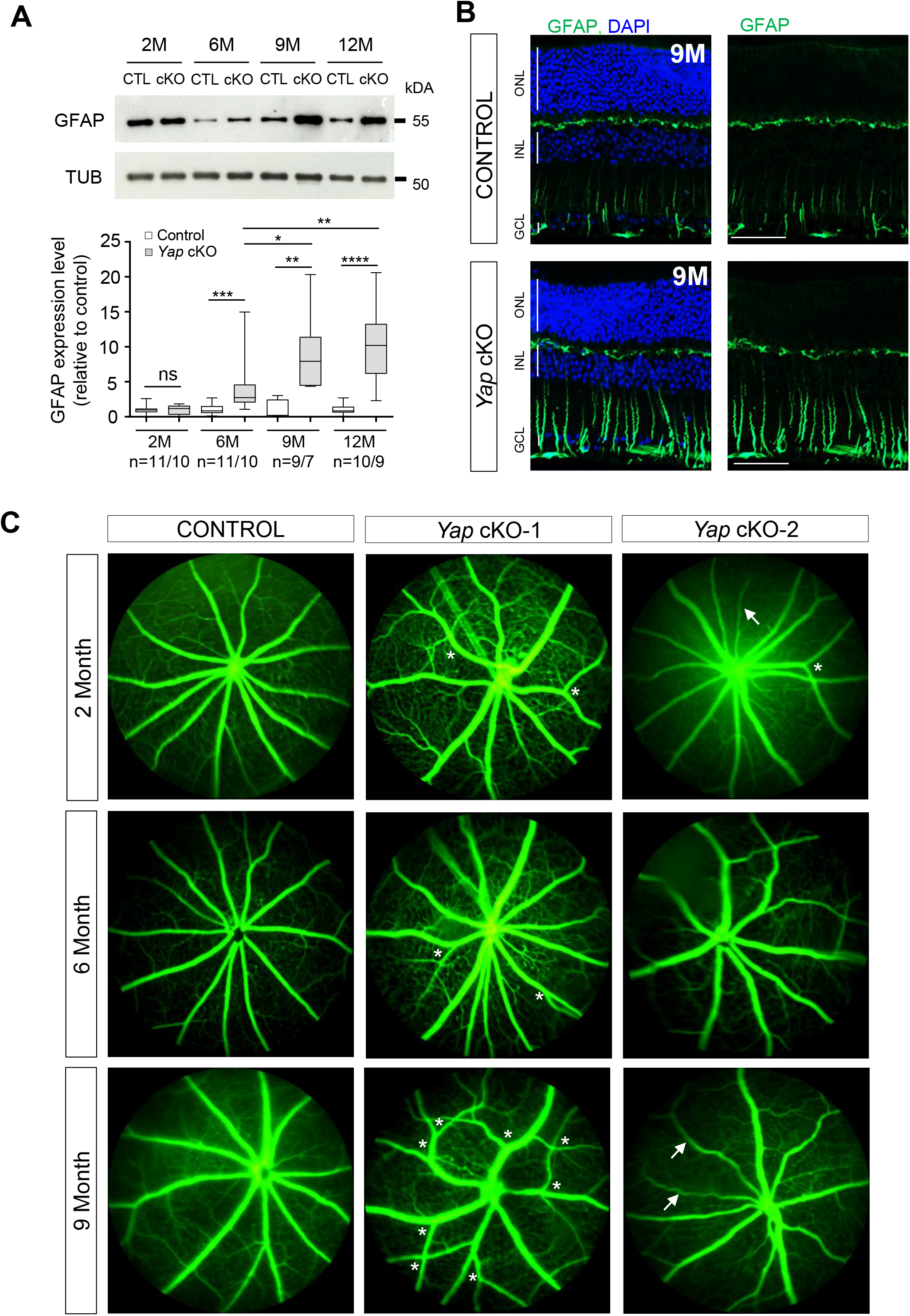
*Yap* cKO retinas exhibit reactive gliosis and vascular defects. (A) Western-blot analysis of GFAP expression in 2-, 6-, 9-, and 12-month-old (2M, 6M, 9M, 12M) control (CTL) and *Yap* cKO retinas. Results in boxplots are expressed after normalization to a-Tubulin (TUB) expression and relative to control. The number of analyzed retinas is indicated below the x-axis. Statistics: Mann-Whitney test, *p≤ 0.05, **p≤ 0.01, *** p≤ 0.001, **** p≤ 0.0001, ns: non-significant. (B) Immunofluorescence analysis of GFAP expression on retinal sections from 9-month-old (9M) control (*Yap*^*flox/flox*^) and *Yap* cKO (*Yap*^*flox/flox*^*;RaxCre*^*ERT2*^) mice. Nuclei were counterstained with DAPI (blue). GCL: ganglion cell layer, INL: inner nuclear layer, ONL: outer nuclear layer. Scale bars: 20 mm. (C) Fluorescein angiography performed in 2-, 6-and 9-month-old (2M, 6M, 9M) control (*Yap*^*flox/flox*^) and *Yap* cKO (*Yap*^*flox/flox*^*;RaxCre*^*ERT2*^) individuals. Note the neovascularization (increased number of branches, white asterisks) and vessel degeneration (white arrows) occurring in *Yap* cKO retinas. Two individuals per age are represented.

Sustained gliosis is suspected to affect the blood-retinal-barrier ^4^, and vascular dysfunction has been shown to participate to RGC degeneration ^25^. We thus assessed retinal blood vessel integrity in *Yap* cKO retinas. Fluorescein angiography revealed retinal vasculature defects starting at 2 months of age (Figure 4C). The phenotype was variable among mice, with defects including excessive branching, tortuous blood vessels, and thin attenuated blood vessels.

These data collectively reveal that *Yap* cKO mice experience vascular defects as early as two months of age, and reactive gliosis starting at six months of age, before the onset of retinal ganglion cell loss.

### Glutamate recycling is impaired in aged Yap cKO retinas

A key regulatory function of Müller cells is to ensure the recycling of glutamate, which is the main excitatory neurotransmitter in the retina ^7^. Control of glutamate levels is crucial for neurotransmission but also to prevent excitotoxicity-dependent RGC death, a phenomenon that has been proposed to participate in glaucoma etiology ^26^. We thus investigated whether RGC loss in *YAP* cKO retinas might result from defective glutamate recycling. We first assessed the glutamate-to-glutamine (Glu/Gln) ratio in 9-month-old retinal extracts by quantitative liquid chromatography coupled to high resolution mass spectrometry. Aged *Yap* cKO retinas harbored a significantly higher Glu/Gln ratio compared to control ones, which results essentially from decreased levels in retinal glutamine content (Figure 5). We then evaluated the expression level of glutamine synthetase (GS), a Müller glia-specific enzyme catalyzing the conversion of glutamate into glutamine ^27^. GS levels in Müller cells were significantly decreased from two months onwards in *Yap* cKO retinas compared to control ones (Figure 6). Consistently with previous data we published ^9^, this was followed in older animals by the downregulation of two other proteins involved in glutamate recycling, namely the inwardly rectifying potassium channel (Kir4.1) ^7^ and the water channel Aquaporin-4 (AQP4) ^28,29^ (Figure 6). Of note, although Kir4.1 was downregulated in Müller cell processes, its expression was increased at their endfeet within the ganglion cell layer.

**Figure 5.**
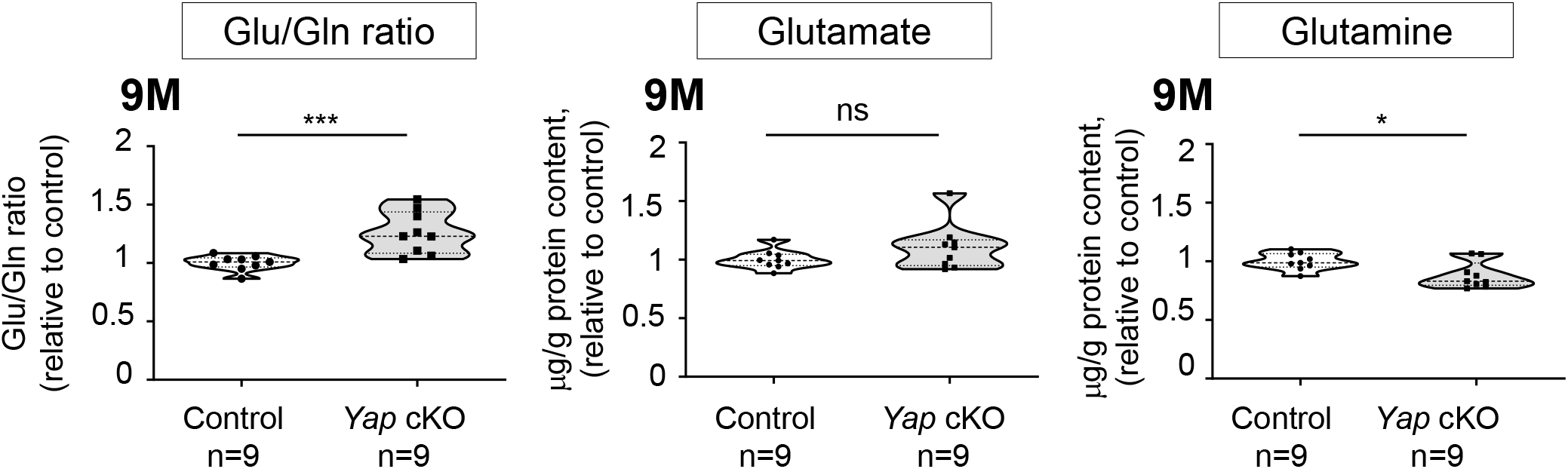
*Yap* cKO retinas show altered glutamate to glutamine ratios. Measurement of the Glutamate to Glutamine ratio (Glu/Gln) and Glutamate or Glutamine content, in 9-month-old (9M) control and *Yap* cKO retinal extracts. The thick dotted line of violin plots, represents the median, and the thin dotted lines the interquartiles. The number of analyzed retinas in each condition is indicated below the x-axis. Statistics: Mann-Whitney test, * p≤ 0.05, *** p≤0.001, ns: non-significant.

**Figure 6.**
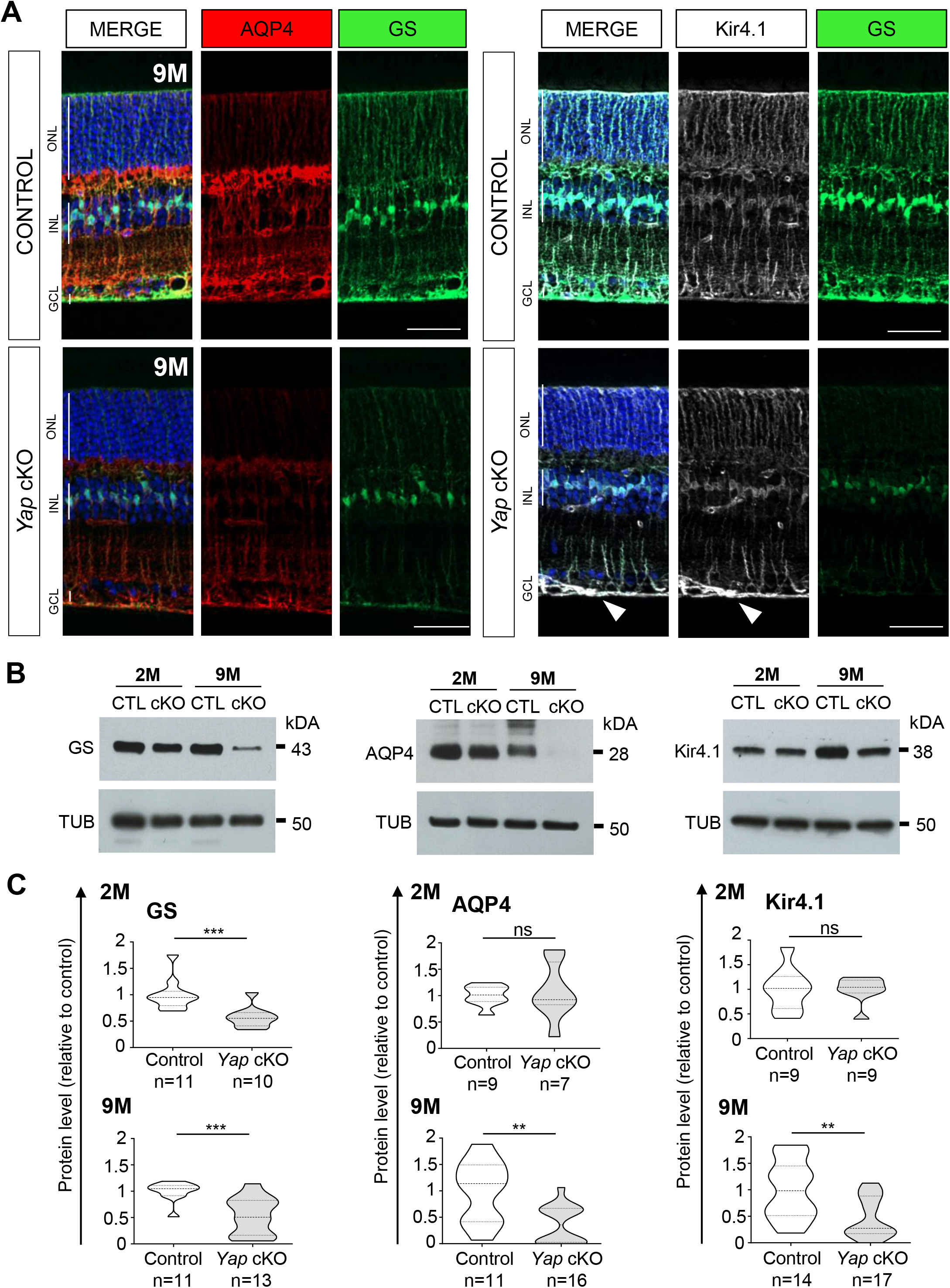
Proteins involved in glutamate recycling are affected in *Yap* cKO Müller cells. Retinal sections from 9-month-old (9M) control (*Yap*^*flox/flox*^) and *Yap* cKO (*Yap*^*flox/flox*^*;RaxCre*^*ERT2*^) retinas, immunostained for glutamine synthetase (GS, green), aquaporin-4 (AQP4, red) and ATP-dependent inwardly rectifying potassium channel 4.1 (Kir4.1, white). Nuclei were counterstained with DAPI. GCL: ganglion cell layer, INL: inner nuclear layer, ONL: outer nuclear layer. The white arrowhead indicates accumulation of Kir4.1 at the endfeet of *Yap* cKO Müller cells. Scale bars: 20 mm. (B) Western-blot analysis of GS, AQP4 and Kir4.1 expression in 2-and 9-month-old (2M, 9M) control (CTL) and *Yap* cKO retinas. (C) Corresponding quantifications. Results in violin-plots are expressed after normalization to a-Tubulin (TUB) expression and relative to control. The thick dotted line represents the median, and the thin dotted lines the interquartiles. The number of analyzed retinas in each condition is indicated below the x-axis indicate. Statistics: Mann-Whitney test, **p≤ 0.01, *** p≤ 0.001, ns: non-significant.

Taken together, these results indicate alteration of glutamate homeostasis in *Yap* cKO retinas, that likely results from defective expression of key Müller glia glutamate recycling proteins.

## DISCUSSION

The mechanisms underlying the pathogenesis of neurodegeneration in glaucoma are still poorly understood. We here document a secondary glaucoma-like phenotype resulting from *Yap* deletion in both the ciliary body NPE and Müller cells, (Figure 7). In the anterior chamber of *Yap* cKO eyes, we found early BaB breakdown and ciliary body collapse resembling the phenotype observed in uveitic patients. In the neuroretina, we highlighted early vascular defects and optic nerve head cupping, and Müller cells dysfunctions that worsen with age (materialized by reactive gliosis and glutamate recycling defects). Ultimately, anterior and posterior chambers dysfunctions led to late-onset of RGCs death. Altogether our results led us to propose that *Yap* cKO mice represent a novel mouse model sharing many important features observed in uveitic glaucomatous patients, that could serve to decipher the molecular mechanisms underlying the disease etiology.

**Figure 7.**
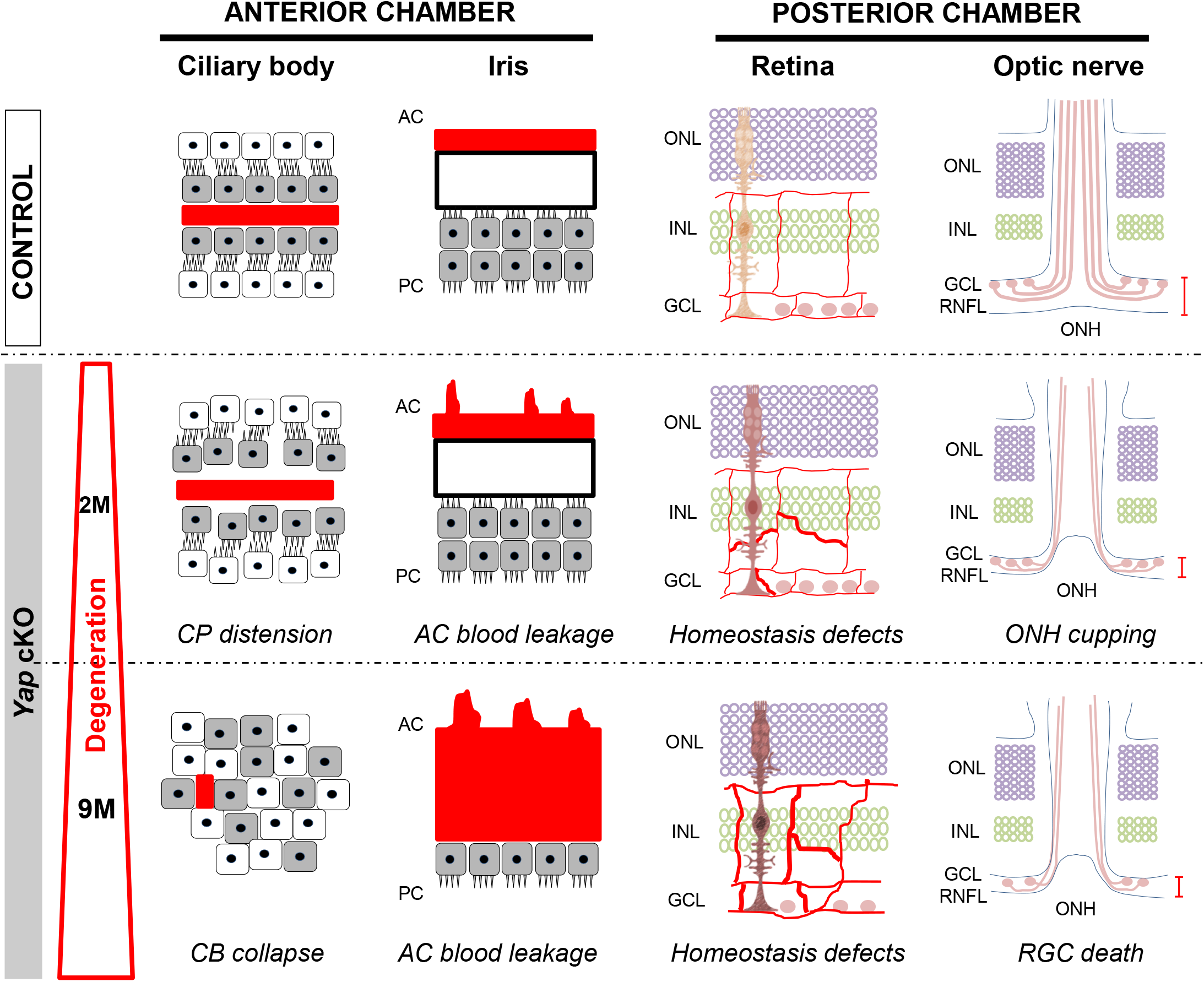
Glaucoma features in *Yap* cKO mice. *Yap* deletion in both Müller glia and NPE cells of the ciliary body leads to defects in homeostatic key functions that eventually result in glaucoma development. Within the anterior chamber, *Yap* cKO mice harbor breakdown of the blood-aqueous-barrier starting by 2 months (2M) at the iris level, and distension of ciliary processes. The ciliary body further degenerates with age until complete collapse at 9 months (9M) of age. At the retinal level, early defects include downregulation of Müller cell proteins including glutamine synthetase and vascular defects accompanied by optic nerve head deformation. This is followed along with aging by reactive gliosis, deregulation of glutamate homeostasis and RGC death. AC: anterior chamber, CB: ciliary body, CP: ciliary processes, GCL: ganglion cell layer, INL: inner nuclear layer, ONH: optic nerve head, ONL: outer nuclear layer, PC: posterior chamber, RGC: retinal ganglion cells, RNFL: retinal nerve fiber layer.

YAP is now recognized as a crucial mediator of eye development, homeostasis and disease ^9,30,31^. Beside its major functions as a cell-autonomous regulator of key cellular processes (proliferation, differentiation, apoptosis…), YAP also acts at the level of the tissue structure ^32^ by bridging extracellular matrix changes to the cell through its mechanosensory activity, and by contributing to the establishment and maintenance of cellular junctions within epithelia ^33^. It is thus important to stress out that the conditional deletion of *Yap* may not only affect NPE and Müller cell transcriptional networks but also result in profound changes within surrounding tissues, through deregulation of local microenvironments.

At the neuroretinal level, *Yap* cKO mice harbor several defects that likely contribute to their glaucoma-like phenotype. In particular, we observed loss of retinal blood vessel integrity as well as Müller cell dysfunctions related to water clearance, ion transport and glutamate recycling. Deregulation of these essential intertwined pathways presumably compromises retinal homeostasis. The water transport is permanently linked to ion fluxes and thereby contributes to neuro-glial communication and neuronal electrical activity. Actually, the coupling of AQP4 with potassium channels like Kir4.1 ensures the establishment of proper K^+^ gradients that are crucial for RGCs physiology ^34,35^. The decreased expression of both proteins observed in aged *Yap* cKO Müller glia may thus contribute to a switch towards a symptomatic retinal phenotype that participates to the onset of RGCs degeneration. Glutamate homeostasis disruption probably represents another important contributor to RGC death. Indeed, under healthy conditions, Müller glia ensure glutamate recycling through its uptake and subsequent transformation into either glutamine, to fuel neurons, or glutathione, the major source of retinal antioxidant ^7^. *Yap* cKO mice present with an altered Glutamate to Glutamine ratio most certainly due to the downregulation of glutamine synthetase expression from two months of age. We did not observe elevated levels of glutamate in retinal extracts, but one cannot exclude that free glutamate might be increased through impaired glutamate reuptake. This function is ensured by the Glutamate-Aspartate Transporter (GLAST) expressed in Müller glial cells. Since its activity is highly dependent on potassium buffering by the Kir4.1 channel ^36^, it might thus be affected by the severely decreased expression of this protein in aged *Yap* cKO retinas. Under this hypothesis, glutamate reuptake may not be optimal and could therefore contribute to local excitotoxicity and eventually RGC death. Of note, such a decreased Müller cell expression of Kir4.1 has been described in glaucomatous patients ^37^, thereby reinforcing the fact that *Yap* cKO mouse phenotype is very close to the human pathology. Finally, increasing evidence support a role of vascular dysfunction in some cases of glaucoma ^25^. In line with this, we observed that RGC loss is not only preceded by reactive gliosis, but also by the appearance of severe retinal vascular defects. These might affect RGC survival by inducing ischemia and subsequent oxidative stress. Altogether, although several questions remain to be addressed, our *Yap* cKO model may help identify the chain of events leading to such neuro-glia vascular unit dysfunctions, and better apprehend how the interplay of different stressors results in RGC degeneration.

In the anterior chamber, the deletion of *Yap* in NPE cells results in ciliary body defects that worsen with age. The early ciliary process distension we observed is indicative of cell-cell contact rigidity loss and is consistent with the known requirement of YAP in cellular junction integrity ^33,38,39^. This is followed by loss of cell polarity and progressive embedment of NPE cells inside the collapsed ciliary body. It was previously shown that gap junction integrity between NPE cells and the pigmented epithelial cells are necessary for the control of IOP ^40^. However, the comparison of *Yap* cKO and control individuals did not reveal significant IOP changes between groups. Of note, except for this aspect, the phenotypic evolution of *Yap* cKO eyes resembles the one observed in DBA/2J mice, a model resulting from mutations in two genes expressed in iris and ciliary body, *Gpnmb* and *Tyrp1* ^21,22^. In both cases, disruption of the iridal BaB precedes the complete collapse of the ciliary body ^21,22^. Since in our model *Yap* is not deleted in the iris, the observed blood leakage is likely an indirect consequence of NPE dysfunction. Based on the intimate topological connection between the iris and ciliary body, an attractive hypothesis would be that mechanical perturbations generated by the disruption of NPE integrity might be transmitted through the iris stroma to the endothelial cell layer, eventually leading to iridal vascular damages.

Glaucoma diseases are multifactorial and therefore involves multiple and intermingled mechanisms. Several preclinical animal models have been developed including induced acute models, spontaneous mutants, and genetically altered models ^41^. Although each of them only reflects one glaucoma subtype with its own etiology, all largely have contributed to major breakthroughs in our understanding of the disease, notably regarding retinal and optic nerve aspects. The most important feature of a reliable model is its reproducibility. In line with this, *Yap* cKO phenotype is fully penetrant with consistent onset and progression among individuals. In addition, *Yap* cKO mice offer the dual benefit to constitute both a uveitis model when considering young individuals, and a glaucoma model developing along with age. To our knowledge, the only other uveitic glaucoma model (mostly used for its glaucoma features) is the DBA/2J model ^22^. It is important to stress out that no uveitis and/or glaucoma model will recapitulate the full spectrum of clinical features in human patients ^42^. To our knowledge, we are the first to report clinical evidence observed in uveitic patients that support the use of *Yap* cKO mice as a relevant model to study the mechanisms underlying the degeneration of the ciliary body. This result is of utmost importance considering the high prevalence (around 20%) of glaucoma complication among uveitic patients ^43^ and the lack of early diagnosis biomarkers. We believe that *Yap* cKO mice could help identifying such biomarkers thereby allowing a better management of glaucomatous patients.

In conclusion, our data bring the first experimental evidence pointing YAP as a key player in glaucoma pathogenesis. They also highlight *Yap* cKO mice as a novel model that might prove useful to elucidate how anterior uveitis may favor glaucoma development and to investigate in the glaucomatous phase how glial and vascular perturbations ultimately result in RGC death. More importantly, this model can offer a new opportunity to test new therapeutic approaches targeting such disease.

## METHODS

### Ethics statement

Ultrasound biomicroscopy was performed in human patients in private practice (ALM-Oftalmolaser, Lisbon, Portugal). Images were anonymized accordingly and adapted with permission of Dr. Helena Prior Filipe and Dr. Maria Sara Patrício. They are part of the Atlas of Ultrasound Biomicroscopy (Design and paginated by Criações Digitais, Lda. Publicated by Théa, Portugal. First edition in May 24th, 2018, https://thea.pt/sites/default/files/documentos/atlas_de_biomicrospia_ultrassonica_ubm_web_low_0.pdf). Updated access November 2020.

All mice were bred and manipulated, in accordance with the European guidelines for the care and ethical use of laboratory animals (Directive 2010/63/EU of the European Parliament and of the Council of 22 September 2010 on the protection of animals used for scientific purposes), and with the Association for Research in Vision and Ophthalmology Statement for the Use of Animals in Ophthalmic and Vision Research. All animal procedures were approved by institutional guidelines, under the license APAFIS#998.

### Animals

*Yap*^*flox/flox* 45^ and *Yap*^*flox/flox*^;*Rax-CreER*^*T2*^ mice ^15^ were kept at 21°C, under a 12-hour light/12-hour dark cycle, with food and water supplied *ad libitum*. Cre recombinase activity was induced through a double intraperitoneal injection of 4-hydroxytamoxifen (4-OHT; 1 mg) at P10 and P14. Of note, the entire progeny was injected with 4-OHT with genotype blinding at the time of injection. Genotyping was performed on tail snip genomic DNA as previously described ^15^. Genotypes and genders of littermates fit with the expected fifty/fifty ratios (Supplementary Table 1). Experiments were performed without gender consideration.

### Retinal dissection

The eyes of sacrificed animals were rapidly enucleated and dissected in Hank’s balanced salt solution (HBSS, Invitrogen). A hole was done with a 33 gauge needle just below the ora serrata. Scissors were inserted horizontally, and a cut was performed all the way below the ora serrata to separate the anterior chamber from the posterior one. The lens was gently removed with forceps and the retina was picked up at the basis of the optic nerve and immediately frozen until further experiments.

### Histology and immunofluorescence

Immunostaining on sections was performed using standard procedures. Briefly, enucleated eyes were fixed in chilled 1X PBS, 4% paraformaldehyde (PFA) for 20 min at 4°C. A hole was done with a 33 gauge needle at the ora serrata and the eyes were fixed another 30 min in PFA. Fixed samples were dehydrated, embedded in paraffin and sectioned with a Microm HM 340E microtome (Thermo Fisher Scientific). Antigen unmasking treatment was done in boiling heat-mediated antigen retrieval buffer (10 mM sodium citrate, pH 6.0) for 20 min. Section permeabilization was performed in PBS 0.3% Triton X-100 for 7 min at RT. All primary and secondary antibodies are listed in Supplementary Table 2. Nuclei were counterstained with DAPI (1mg/ml, Thermo Fisher Scientific) for 10 min.

For NEFL immunostaining on flat-mounted retinas, retinas were dissected in HBSS and all the vitreous was removed in order to optimize antibody binding. The retinas were then fixed in 4% PFA for 20 min, washed with PBS and permeabilized/blocked in DAKO diluent (DAKO) containing 0.5% Triton X-100 for 7 minutes at RT. The same solution was used for incubation in primary and secondary antibodies. Four radial incisions were then made in the retina to create a petal shape and allow flat mounting with ganglion cells facing up.

### Imaging

Fluorescence and brightfield images of paraffin-embedded eye sections were acquired using an ApoTome-equipped AxioImager.M2 microscope. Image processing was performed using Zen (*Zeiss*), and ImageJ softwares ^45^. The same magnification, laser intensity, gain and offset settings were used across animals for any given marker. Ultrasound biomicroscopy images were acquired using a Reflex UBM (Reichert Technologies).

### Western-blot

Western-blot was performed on proteins extracted from a single retina. Between seven and sixteen individuals were tested per condition. Harvested retinas were snap frozen before lysis in 80 µL P300 buffer (20 mM Na2HPO4; 250 mM NaCl; 30 mM NaPPi; 0.1% Nonidet P-40; 5 mM EDTA; 5mM DTT) and protease inhibitor cocktail (Sigma-Aldrich). Protein content was measured using the Lowry protein assay kit (DC Protein Assay; Bio-Rad). Homogenates were sonicated and centrifuged for 15 min at 5000 g, and then 10 μg of the supernatant were subjected to SDS-PAGE, as previously described ^11^. Western blots were then conducted using standard procedures. Primary and secondary antibodies are listed in Supplementary Table 2. Antibody binding was revealed by the Enhanced Chemiluminescence System (Life Technologies) on X-Ray film (Sigma-Aldrich). Each sample was probed once with anti-α-tubulin antibody for normalization. Quantification was done using ImageJ software ^45^.

### Measurement of the Glutamate-to-Glutamine ratio by liquid chromatography coupled to high-resolution mass spectrometry (LC-HRMS)

Retinas were resuspended in 170 µL of ultrapure water, and then sonicated 2 times for 10 s using a sonication probe (vibra cell, Bioblock Scientific). At this step, 20 µL of each sample were withdrawn for total protein concentration determination (Pierce BCA Protein Assay Kit, Thermo Fisher Scientific). Then, we added to the remaining lysate 350 µL of methanol and 10 μL of an internal standard mixture containing 50 µg/mL of ^13^C5-glutamine (Eurisotop) and ^13^C5-glutamic acid (Sigma). Following a 1.5 hour incubation on ice, cell debris were removed by centrifugation for 15 min at 4 °C and 20,000 g. The resulting metabolic extracts were dried under a stream of nitrogen using a TurboVap instrument (Thermo Fisher Scientific) and stored at −80 °C until analysis. Dried extracts were dissolved in 30 μL of mobile phase A (see below), sonicated, vortexed, and mixed with 70 µL of mobile phase B, sonicated and vortexed again before a final centrifugation step at 4 °C for 5 min at 20,000 g. The resulting supernatants were then transferred into vials for analysis by LC-HRMS.

LC-HRMS quantification of glutamine and glutamic acid was performed using a Dionex Ultimate chromatographic system (Thermo Fisher Scientific) coupled to a Q-Exactive Plus mass spectrometer (Thermo Fisher Scientific) equipped with an electrospray ion source. The mass spectrometer was externally calibrated before each analysis according to the manufacturer instructions. Chromatographic separation was performed on a Sequant ZIC-pHILIC column (5 μm, 2.1 × 150 mm; Merck) maintained at 45°C. Mobile phase A consisted in an aqueous buffer of 10 mM of ammonium acetate, while mobile phase B was made of 100% acetonitrile. Chromatographic elution was achieved at a flow rate of 200 μL/min. After injection of 10 μL of sample, elution started with an isocratic step of 2 min at 30% A, followed by a linear gradient from 30 to 60% of phase A during 5 min. The chromatographic system was then rinsed for 5 min at 0% B, and the run ended with an equilibration step of 8 min. The column effluent was directly introduced into the electrospray source of the mass spectrometer, and analyses were performed in the negative ion mode. Source parameters were as follows: capillary voltage, -2.5kV; capillary temperature, 350°C; sheath gas and auxiliary gas pressures set at 35 and 10 arbitrary units, respectively. The detection was performed from m/z 50 to 600 at a resolution of 70,000 (full width at half-maximum at m/z 200) using an AGC target of 1e6 and a maximum injection time of 250 ms. Glutamine and glutamic acid were detected as deprotonated [M-H]-species at m/z 145.0618 and m/z 146.0458, and elute at 4.7 and 4.8 min, respectively. Both entities were quantified by isotope dilution using their corresponding ^13^C-labeled homologues as internal standards and concentrations expressed as µg of glutamine or glutamic acid per gram of total proteins.

### Measurement of intraocular pressure (IOP)

IOP was measured in both eyes with a rebound tonometer (Tono-Lab, Tiolat, OY) ^46^ on 6-and 9-month-old awaken mice. At each time-point, 18 consecutive readings were made for each eye and averaged. To limit fluctuations of the IOP due to the circadian rhythm, we always measured IOP in the morning ^46^.

### Retinal and anterior chamber angiography

Mice were anesthetized with intraperitoneal injection of ketamine (90 mg/ kg, Merial) and xylazine (8 mg/kg, Bayer). Fluorescein angiography was performed by injecting 150 µL dextran-conjugated fluorescein (25mg/ml final, molecular weight [MW] 10,000) into the vein tail of anesthetized mice. Images were taken for up to 5 minutes post-injection after pupil dilation by topical application of tropicamide (0.5%) and phenylephrine (2.5%) for retinal images, or without pupil dilation for pupillary dilation for anterior chamber images, using a Micron IV retinal imaging microscope (Phoenix Research Labs).

### Optic coherence tomography (OCT)

Mice were anesthetized with intraperitoneal injection of ketamine (90 mg/ kg, Merial) and xylazine (8 mg/kg, Bayer). Pupil were dilated by topical application of tropicamide (0.5%) and phenylephrine (2.5%). OCT was performed using the R2200 Spectral Domain OCT system Imaging System (Bioptigen Inc.). For optic nerve head imaging, a general mouse OCT lens was used with reference arm set to 1020. A rectangular volume analysis was performed, using 33 consecutive B-scans lines, each one composed of 1000 A-scans and three frames. The volume diameter was 1.4 mm.

### Ultrasound biomicroscopy (UBM)

To visualize the anterior chamber angle including the ciliary body and ciliary processes, UBM examination was performed in patients in supine position, by one operator, in mesopic conditions, after topical anesthesia with oxybuprocaine hydrochloride (Anestocil®) with a Reflex UBM (Reichert Technologies). The plastic shell was preferred to prevent globe distortion, and immersion was accomplished with distilled water and Hypromellose with Carbomer 980 (Genteal Eye gel) applied along the shell’s interior rim to keep the bath sealed. Patients were instructed to gaze straight ahead at one point in the ceiling to acquire as close as possible the axial sections. For transverse sections, patients were asked to gaze at the opposite direction of the meridian under observation, and the ultrasound probe was placed over the latter with its marker towards up or nasally for a transverse swipe.

### Statistics

All data were analyzed using GraphPad Prism version 8.3 for Windows (GraphPad Software, La Jolla California USA; www.graphpad.com). Comparisons involving two unpaired groups were analyzed using the non-parametric two-tailed Mann-Whitney test. A *P* value <0.05 was set as the basis for rejecting the null hypothesis. Sample sizing is depicted in the figure legends.

## Author contributions

Experiments were performed by JB except angiography which was performed by EG, OCT which was performed by SL, glutamate/glutamine ratio determination which was performed by FF, FC, and ECV (MetaboHub platform), IOP measurements which were performed by JB and AH. HPF performed the UBM examination in human patients. JB conceived the experiments. JB, MP, JR, and ML analyzed the data and wrote the manuscript.

## Acknowledgements

This work was supported by the Commissariat à l’Energie Atomique et aux Energies Alternatives and the MetaboHUB infrastructure (ANR-11-INBS-0010 grant). We thank Rémi Souillé for his technical assistance. This research was supported by grants to MP from Association Retina France, Fondation Valentin Haüy and UNADEV.

---

**Supplemental table 1.** Genotype and sex repartition in the mouse colony.

| Category | Numbers |
| --- | --- |
| Mice* | 112 |
| Control | 50 |
| *Yap* cKO | 62 |
| Females | 57 |
| Males | 55 |
| Control females | 27 |
| *Yap* cKO females | 30 |
| Control males | 23 |
| *Yap* cKO males | 32 |

\* Age between 2- and 9-months

**Supplemental Table 2.** List of antibodies.

| Antigen | Host | Supplier | Reference | Dilution |  |
| --- | --- | --- | --- | --- | --- |
|  |  |  |  | Dilution (IHC) | (WB) |
| Primary antibodies |  |  |  |  |  |
| Aquaporin-4 | Rabbit | Alomone Labs | AQP-004 | 1:1,000 | 1:750 |
| Brn3a | Mouse | Santa Cruz | sc-8429 | 1:100 |  |
| GFAP | Rabbit | Dako | Z0334 | 1:500 |  |
| GFAP | Goat | Abcam | ab53554 |  | 1:1,500 |
| Glutamine synthetase | Mouse | Abcam | ab64613 | 1:3,000 | 1:3,000 |
| Kir4.1 | Rabbit | Alomone Labs | APC-035 | 1:1000 | 1:750 |
| NEFL | Mouse | EMD Millipore | MAB1615 | 1:100 |  |
| Sox9 | Rabbit | EMD Millipore | AB5535 | 1:300 |  |
| α-tubulin | Mouse | Sigma-Aldrich | T9026 |  | 1:5,000 |
| Albumin-FITC | Goat | Bethyl Laboratories | A90-234F | 1:300 |  |
| Secondary antibodies |  |  |  |  |  |
| Alexa 488 anti-rabbit | donkey | Thermo Fisher Scientific | A21206 | 1:500 |  |
| Alexa 555 anti-rabbit | donkey | Thermo Fisher Scientific | A31572 | 1:500 |  |
| Alexa 488 anti-mouse IgG1 | goat | Thermo Fisher Scientific | A21121 | 1:500 |  |
| Alexa 555 anti-mouse IgG1 | goat | Thermo Fisher Scientific | A21127 | 1:500 |  |
| Alexa 555 anti-mouse IgG2a | goat | Thermo Fisher Scientific | A21127 | 1:500 |  |
| Alexa 555 anti-mouse IgG2b | goat | Thermo Fisher Scientific | A21147 | 1:500 |  |
| Alexa 488 anti-mouse | goat | Thermo Fisher Scientific | A11001 | 1:500 |  |
| Anti-rabbit HRP | Goat | GE Healthcare | NA-934V |  | 1:5,000 |
| Anti-mouse HRP | Goat | Sigma-Aldrich | A4416 |  | 1:5,000 |
| Alexa conjugated lectin |  |  |  |  |  |
| Isolectin-B4 alexa 594 conjugate | Griffonia simplicifolia | Thermo Fisher Scientific | I21413 | 1:300 |  |

